# A Robust Spike Sorting Method based on the Joint Optimization of Linear Discrimination Analysis and Density Peaks

**DOI:** 10.1101/2022.02.10.479846

**Authors:** Yiwei Zhang, Jiawei Han, Tengjun Liu, Zelan Yang, Weidong Chen, Shaomin Zhang

## Abstract

**Objective:** Spike sorting is a fundamental step in extracting single-unit activity from neural ensemble recordings, which play an important role in basic neuroscience and neurotechnologies. A few algorithms have been applied in spike sorting. However, when noise level or waveform similarity becomes relatively high, their robustness still faces a big challenge.

**Approach:** In this study, we propose a spike sorting method combining Linear Discriminant Analysis (LDA) and Density Peaks (DP) for feature extraction and clustering. Relying on the joint optimization of LDA and DP: DP provides more accurate classification labels for LDA, LDA extracts more discriminative features to cluster for DP, and the algorithm achieves high performance after iteration. We first compared the proposed LDA-DP algorithm with several algorithms on one publicly available simulated dataset and one real rodent neural dataset with different noise levels. We further demonstrated the performance of the LDA-DP method on a real neural dataset from non-human primates with more complex distribution characteristics.

**Main results:** The results show that our LDA-DP algorithm extracts a more discriminative feature subspace and achieves better cluster quality than previously established methods in both simulated and real data. Especially in the neural recordings with high noise levels or waveform similarity, the LDA-DP still yields a robust performance with automatic detection of the number of clusters.

**Significance:** The proposed LDA-DP algorithm achieved high sorting accuracy and robustness to noise, which offers a promising tool for spike sorting and facilitates the following analysis of neural population activity.

## Introduction

The development of neuroscience has put forward high requirements for analyzing neural activity at both single neuron [1–4] and population levels [5–7]. The basis of neural data analyses is the correct assignment of each detected spike to the appropriate units, a process called spike sorting [8–11]. Spike sorting methods will encounter difficulties in the face of noise and perturbation. Since the algorithms commonly fall into two processes [12–13], feature extraction and clustering, an outstanding spike sorting algorithm needs to be highly robust in the feature extraction and clustering process.

For feature extraction methods, the extracted features are descriptions of spikes in low-dimensional space. An appropriate feature extraction method can reduce data dimensions while ensuring the degree of differentiation [1]. Currently, extracting the geometric features of waveforms is the simplest way, including peak-to-peak value, width, zero-crossing feature, etc. [14]. Although this method is easy to operate with extremely low complexity, it has a low degree of differentiation for similar spikes and is highly sensitive to noise [12–13]. First and Second Derivative Extrema (FSDE) calculates the first and second derivative extrema of spike waveform, which is relatively simple and has certain robustness to noise [15]. Other methods like Principal Components Analysis (PCA) [13,16] and Discrete Wavelet Transform (DWT) [17–19] have certain robustness to noise. In recent years, deep learning methods have been proposed, such as the 1D-CNNs [20] and the Autoencoder (AE) [21]. However, the deep learning methods are limited in practical use because of their high computational complexity and high demands on the training set.

So far, many previous methods tend to be perturbed by noise or the complexity of the data. They cannot effectively extract the features with high differentiation, resulting in poor effect in the subsequent clustering, especially in the case of high noise level and data similarity. For solving this problem, some studies improved the robustness by using supervised feature extraction and clustering iteration to get the optimal subspace with strong clustering discrimination [22–25].

Clustering algorithms are also developing with the update of data analysis methods. Early on, the commonly used method was manually [26] segmenting clusters. When more channels come, the workload of operations becomes higher, so it is less used later. K-means (Km) [13] is a widely used clustering method because it is simple to calculate. However, it requires users to determine the number of clusters in advance. Thus, it is sensitive to the initial parameters and lacks robustness [27–28]. Some distribution-based methods, such as Bayesian Clustering [13] and Gaussian Mixture Model [29–31], represent the data with Gaussian-distribution assumptions. Some methods based on neighboring relations can avoid assumptions, for example, the Superparamagnetic Clustering [19,32]. In addition, Neural Networks [33], T-distribution [34], Hierarchical Clustering [35], and Support Vector Machines [36,37] are also used in spike sorting.

For supervised feature extraction methods, the clustering method has a powerful influence on the performance of feature extraction and further affects the performance of the whole algorithm. Ding et al. proposed the LDA-Km algorithm, which used K-means to obtain classification labels and LDA to find the feature space based on the labels and then continuously iterated the two algorithms to convergence [24–25]. Keshtkaran et al. introduced a Gaussian Mixture Model (GMM) based on LDA-Km and put forward the LDA-GMM algorithm [22], which had high accuracy and strong robustness against noise and outliers. LDA-GMM needs to iterate several times by changing the initial value of important parameters (such as the initial projection matrix) to obtain the optimal result. Its operation also calls LDA-Km which brings additional computation complexity. Recently, the concept of joint optimization of feature extraction and clustering has been adopted to construct a unified optimization model of PCA and Km-like procedures [38], which integrates the feature extraction and clustering steps for spike sorting.

Inspired by these efforts, we proposed a framework that integrates the supervised feature extraction and the clustering to make them benefit each other. Thus, a remarkable clustering method is also crucial. Density Peaks (DP) proposed by Rodriguez et al. define the cluster centers as local maxima in the density of data points [39]. This algorithm does not assume the data distribution and can well adapt to the nonspherical distribution, which is more applicable to the complex distribution of spikes in vivo, making it a win-win for robustness and computation cost. Therefore, this paper integrates the LDA and DP as a joint optimization model LDA-DP for spike sorting.

## Methods

### 1. An overview of the LDA-DP algorithm

This study proposed a spike sorting algorithm combining LDA with DP. LDA is a supervised machine learning method that requires prior information about cluster labels. The data is initially projected into an initial subspace and then clustered by Density Peaks to obtain cluster labels. Thus, we need an initial projection matrix *W*. As summarized in Algorithm 1, the projection matrix *W* is initialized by executing PCA on spike matrix X, cutting the first *d* coefficients, and assigning it to *W*. We chose *d*=3 for overall consideration to maintain performance and computation complexity, in line with the other feature extraction methods compared in this study. In each iteration, the algorithm updates a clustering result *L*. When the updated *L* is relatively consistent with the result in the previous iteration (*L_pre*) or the number of iterations reaches the upper limit *MaxIte*, the iteration ends. The minimum number of iterations *MinIte* ensures that the algorithm iterates adequately. The suggested value for minimum iteration *MinIte* is 5 and maximum iteration *MaxIte* is 50. Finally, in the last step of the algorithm, the similar clusters are merged and we obtain the sorting result *L_merge_*.

#### Algorithm 1. Spike sorting based on LDA-DP

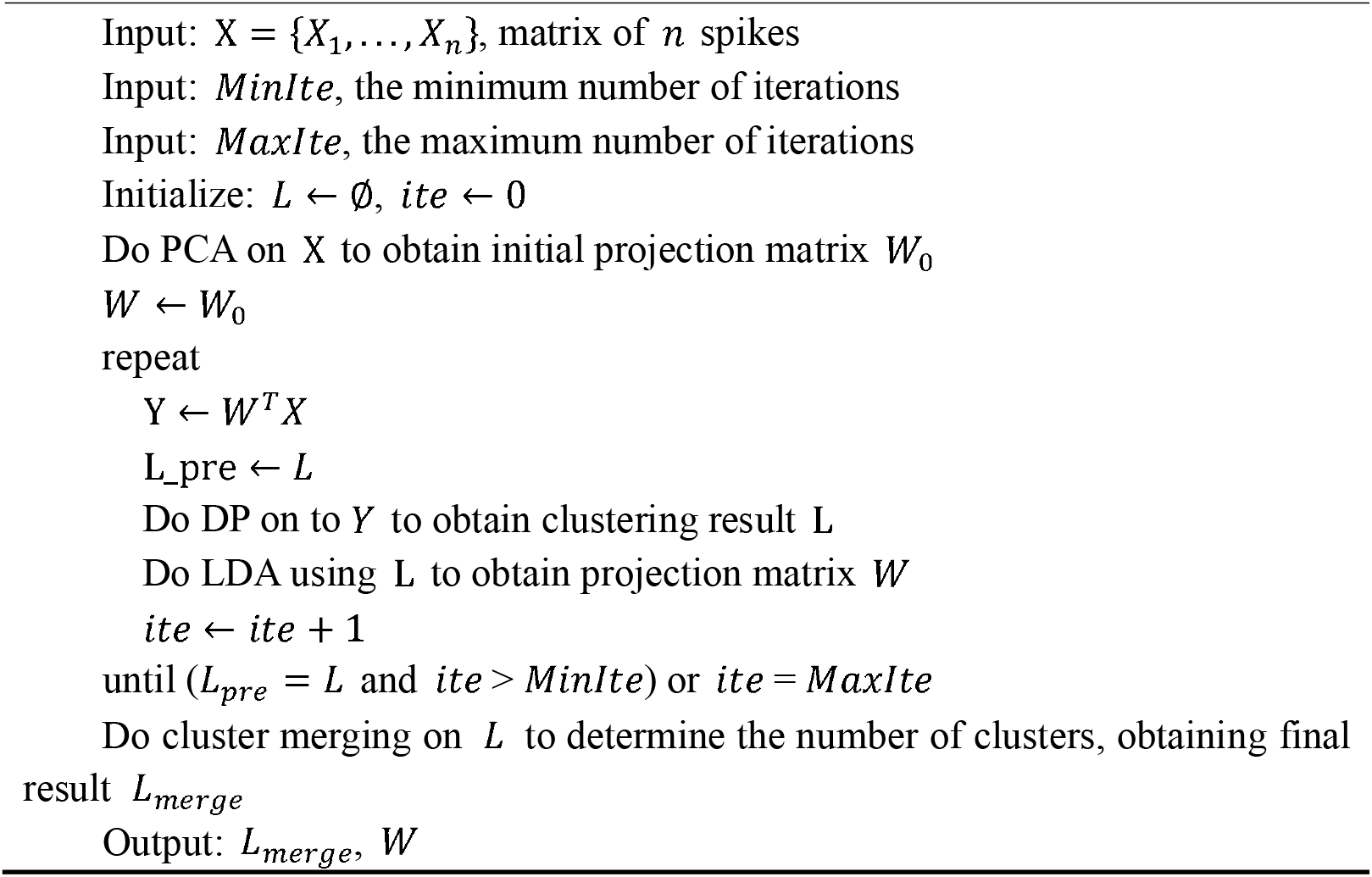

### 2. Discriminative feature extraction using Linear Discriminant Analysis

Linear Discriminant Analysis (LDA), also known as “Fisher Discriminant Analysis”, is a linear learning method proposed by Fisher [23]. LDA is a supervised machine learning method that finds an optimal feature space, where the intra-class scatters are relatively small and inter-class scatters are relatively large.

For a multi-cluster dataset, the quality of clusters can be measured by the intra-class scatter metric *S_w_* and the inter-class scatter metric *S_b_*, as shown in Formula (2.1) and (2.2):

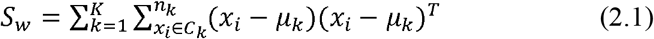

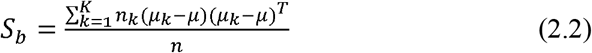

*x_i_* denotes the *i* th data point in the *k* th cluster *C_k_, μ_k_* denotes the mean value of data points in *C_k_, n_k_* denotes the number of data points in *C_k_*,μ denotes the mean value of all data points, and *n* denotes the total number of data points.

To calculate the projection matrix *W*, LDA performs optimization by maximizing objective function *J* (Formula (2.3)).

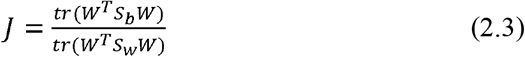

Then data points can be projected to a *d*-dimensional subspace which captures discriminative features by the obtained projection matrix *W*. In this study, *d* was fixed to 3 by default.

### 3. Clustering features based on Density Peaks

The groundwork of the Density Peaks Algorithm (DP) [39] is very simple. For each data point, two parameters are calculated, the local density *ρ* of the point and the minimum distance *δ* between the current point and the data point with a larger local density. The DP algorithm assumes that if a point is a cluster center, it will satisfy two conditions: (1) its local density *ρ* is high; (2) it is far away from another point that has a larger local density. That is, the center of the cluster is large for both *ρ* and *δ*. After the cluster centers are identified, the remaining data points are allocated according to the following principle: each point falls into the same cluster with its nearest neighbor point *n_up* who has a higher local density.

In this study, the Gaussian kernel is adopted to calculate local density. Local density *ρ* of the *i*th point is shown in Formula (2.4):

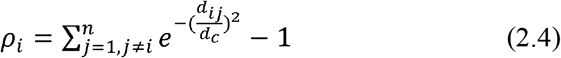

*d_i,j_* donates the euclidean distance between the sample *y_i_* and *y_j_*, as is shown in Formula (2.5):

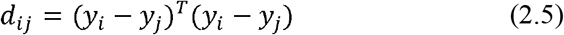

*d_c_* donates the cutoff distance. In this study, we defined cutoff distance by selecting a value in ascending sorted sample distances *d_sort*:

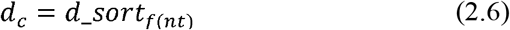

*t* is the cutoff distance index, and as a rule of thumb, it generally ranges from 0.01 to 0.02, and *f*(·) denotes the rounding function.

The minimum distance *δ* and the nearest neighbor point *n_up* is calculated in Formula (2.7) and (2.8) where *ρ_max_* denotes the maximum local density:

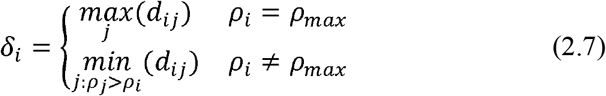

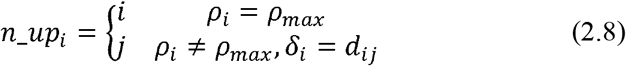

To automate [39] the search for the cluster centers where both *ρ* and *δ* are large, we creatively defined the DP index *λ* as the product of *ρ* and *δ*, as shown in Formula (2.9).

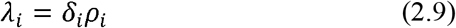

The algorithm selects *K* data points with the largest DP index as the clustering centers. If the data is randomly distributed, then the distribution of *λ* is in line with the monotonically decreasing power function, and the DP index of the cluster centers is significantly higher, which makes it feasible to select cluster centers according to the DP index *λ*. As iteration times increase, the DP index difference between the center and the non-center increases. As the *K* value is determined, the method can be automated.Since *K* denotes the initial number of clusters, its default value in this study is 4.

For dataset *X* which contains *n* data points 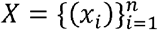, giving cutoff distance index *t* and the initial number of clusters *K*, the flow of the DP clustering algorithm is as follows:

step1: Calculate the distance between every two data points *d_ij_*(*i,j* = 1,2,… *n, i* < *j*), as shown in Formula (2.5).
step2: Calculate the cutoff distance *d_c_* as shown in Formula (2.6).
step3: For each data point, calculate the local density *ρ*, the minimum distance *δ*, and nearest neighbor point *n_up*, as shown in Formula (2.4) (2.7) and (2.8).
step4: For each data point, calculate the DP index *λ*, as shown in Formula (2.9).
step5: Select cluster centers: the point with the largest *λ* is the center of Cluster 1,the point with the second-largest *λ* is the center of Cluster 2, and so on to get *K* centers.
step6: Classify non-center points: rank the non-center points in descending order according to their local density *ρ*, traversing each non-center point, and then the label *L_i_* of the *i* th point is calculated as Formula (2.10):

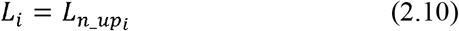

where *L_n_uPi_* denotes the cluster label of the nearest neighbor point of the *i* th point.

Here, we demonstrated the process using the testing set C1_005 (see Evaluation). In the feature extraction step, LDA finds out the feature subspace with the optimal clustering discrimination through continuous iteration (Figure 1a). For each data point, our algorithm calculates its local density (*ρ*) and the minimum distance (*δ*). As described in the previous section, cluster centers are the points whose *ρ* and *δ* are relatively large. DP screens the cluster centers using a previously defined DP index *λ* that is the product of *ρ* and *δ*. Figure 1b shows a schematic diagram where *ρ* and *δ* are set as the horizontal and vertical axes in the case of screening three cluster centers. The screened centers are the three points with the largest *λ*. They are circled in three colors corresponding to three clusters. The defined DP index is competent for center point screening for the screened points all have a large *ρ* and *δ* value. As a result, Density Peaks clustering obtains the three clusters (Figure 1c), and Figure 1d shows the waveforms of each cluster center.

**Figure 1.**
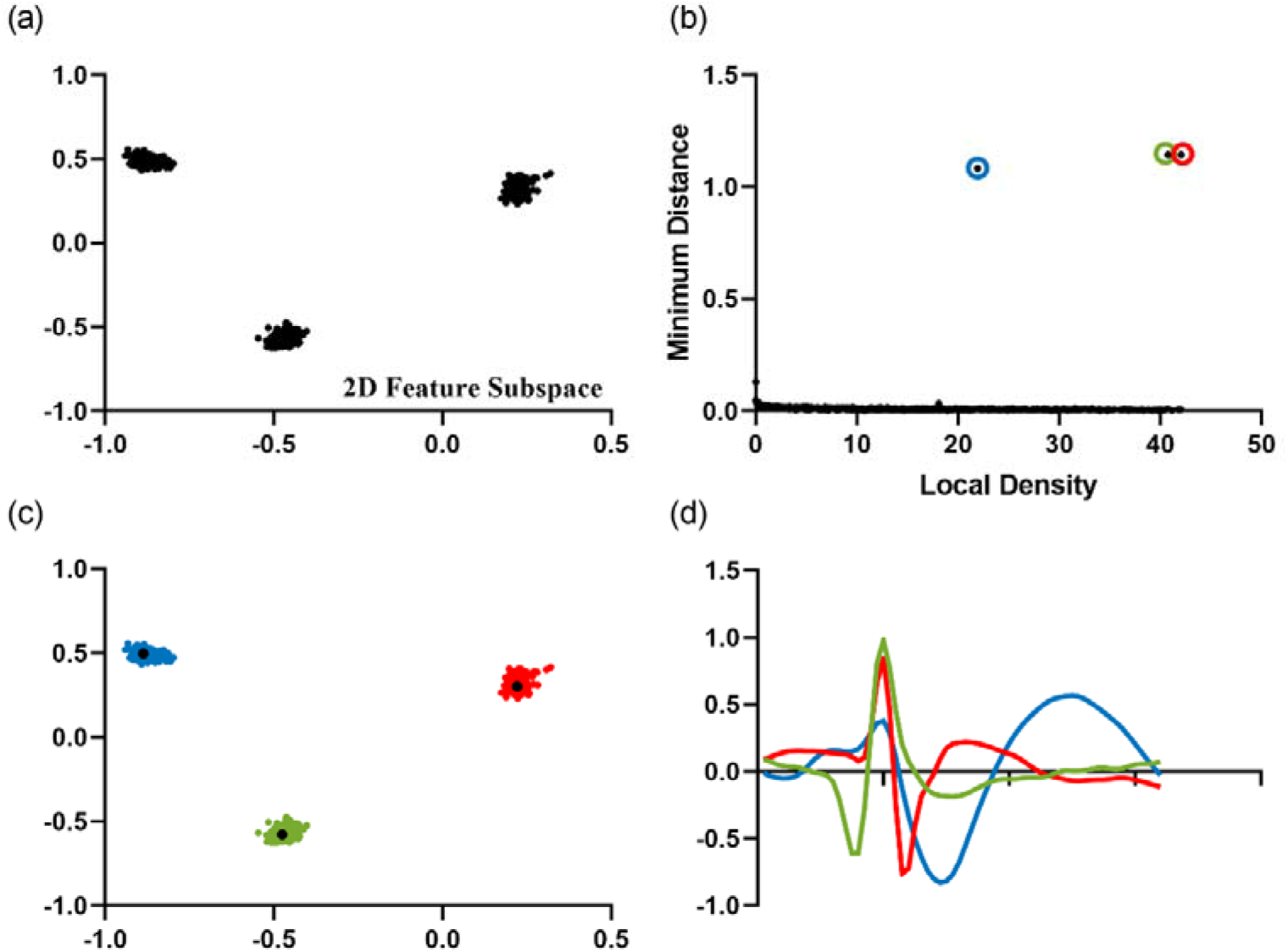
Running scenario of the DP clustering stage using Dataset C1_005. (a) Data points projected in a two-dimensional feature subspace after LDA. This subspace is considered to be optimal for the within-class scatters are relatively small and intra-class scatters are relatively large. (b) Schematic diagram with the local density *ρ* and the minimum distance *δ* as the horizontal and vertical axes. The points circled in three colors are the screened cluster centers. (c) Scatter plot of 3 clusters obtained by DP. The three colors of red, green and blue respectively represent the clustering results of cluster 1, cluster 2 and cluster 3. The black dots represent the cluster centers. (d) The waveforms of three cluster centers.

### 4. Automatic detection of the number of clusters

The last step of the algorithm is a cluster merging step, through which the number of clusters will be determined. The purpose of cluster merging is to avoid similar clusters being over-split. After the merging step, the number of clusters is determined automatically. The number of clusters is a critical parameter required by many spike sorting algorithms. However, manually setting the number of clusters in advance relies heavily on the experience of operators and may cause problems in practice. Thus, a merging step is crucial to automatically determine the number of clusters, in order to reduce the workload and artificial error of manual operation. The cluster merging finds similar clusters, combines them and repeats the process. Here, a threshold is used to end the cluster merging process. Once the similarity between the most similar clusters goes below the threshold, the merging is stopped, and thus the number of clusters is automatically determined.

The similarity between clusters can be measured in several ways. Common distance metrics include the Minkowski distance, the cosine distance and the inner product distance [40]. According to the Davis-Bouldin Index (DBI) [40–43], we defined cluster similarity as the ratio *R* of the compactness *CP* and the separation *SP*.

Intra-class distance is a parameter to evaluate the internal compactness of a cluster. Thus, the compactness *CP* can be calculated by (2.11).

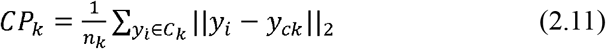

*CP_k_* denotes the within-class distance of the cluster *C_k_, y_i_* denotes the *i* th data point in the *k*th cluster *C_k_, y_ck_* denotes the center of *C_k_*.

Inter-class distance is a parameter to evaluate the separation of clusters. Thus, the separation *SP* can be calculated by (2.12).

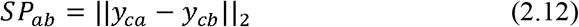

*SP_ab_* denotes the inter-class distance between the cluster *C_a_* and *C_b_*, and *y_ca_* and *y_cb_* denotes the center of the cluster *C_a_* and *C_b_*, respectively.

Similarity metric *R* is shown in Formula (2.13)

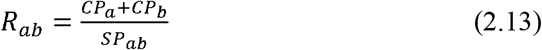

If two clusters have high similarity, the two clusters are merged. We set the threshold *R_th_* as a proportional function of the mean value of *R*, as is shown in Formula (2.14)

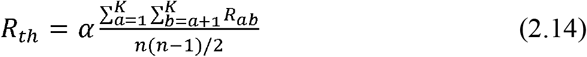

*α* denotes the threshold coefficient. The threshold should be significantly higher than the mean value, and as a rule of thumb, *α* is above 1.4.

The flow of the cluster merging process is as follows:

step1: Calculate the compactness *CP* for each cluster, as shown in Formula (2.11)
step2: Calculate the separation *SP_ab_* (*a, b*= 1,2,… *K, a* < *b*) for every two clusters, as shown in Formula (2.12)
step3: Calculate similarity metric *R_ab_* for for every two clusters, as shown in Formula (2.13)
step4: Calculate the threshold *R_th_*, as shown in Formula (2.14)
step5: Find the maximum similarity *R_ab_*. If *R_ab_ > R_th_*, merge cluster *a* and cluster *b*, set the center of cluster *a* as the new center, *K* = *K* − 1, return to step 1; Otherwise, stop merging.

Notably, the threshold coefficient *α* largely affects the merging results. Figure 2a shows the influence of the threshold coefficient α on algorithm performance (accuracy) on Dataset A (see Evaluation). The mean accuracy reaches the highest when *α* = 1.6. Therefore, we found an appropriate value of α = 1.6 to make the algorithm achieve a general optimal performance on all datasets. In subsequent evaluation, we fixed α as 1.6.

**Figure 2.**
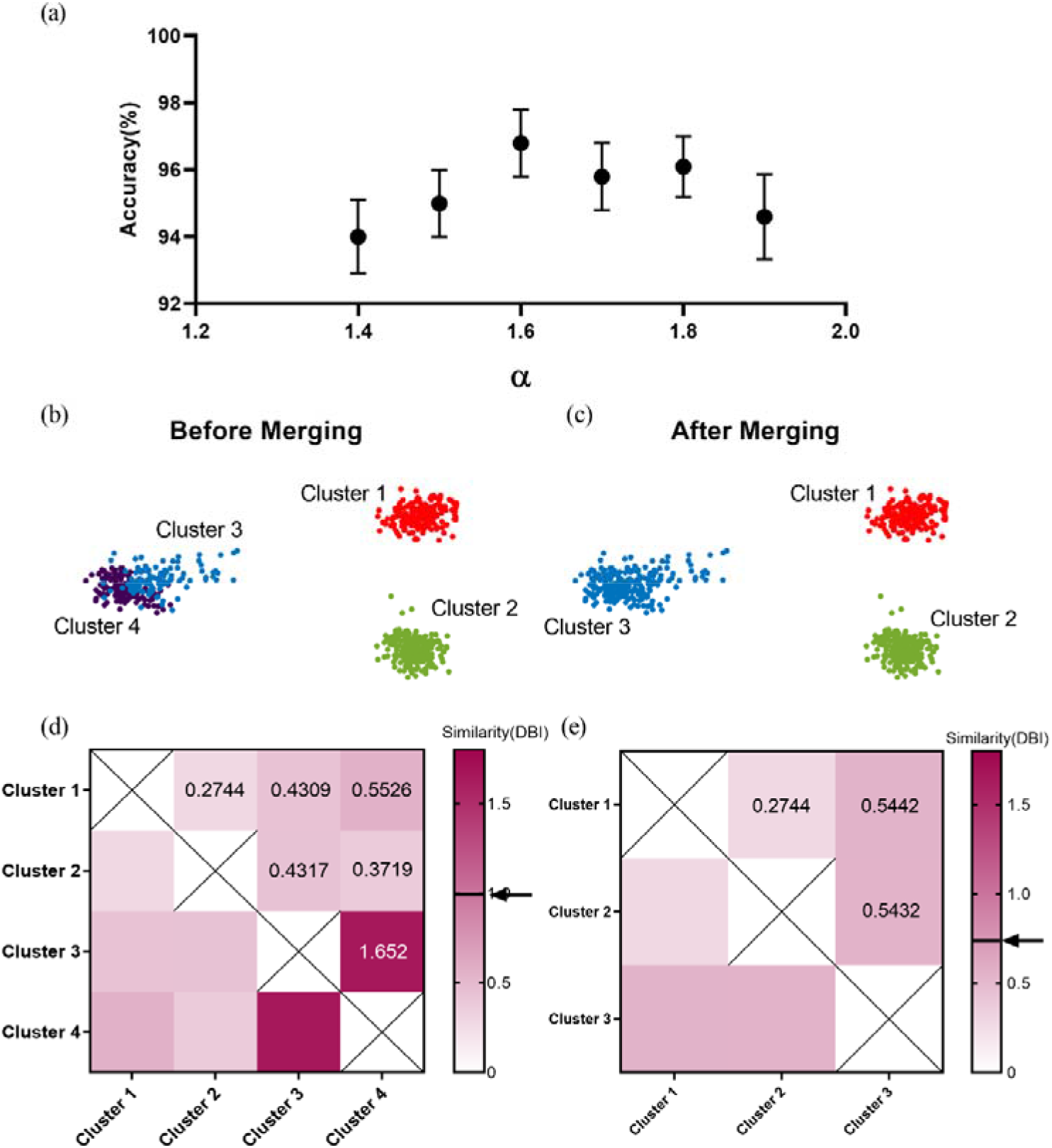
An illustration of determining the number of clusters. (a) Accuracy on all testing sets in dataset A versus threshold coefficient α. The error bars represent the standard error of the mean. (b) (c) The results of classification before and after merging. Before merging, data points are clustered into four clusters (denoted by four colors: red, violet, green and blue), while after merging, we obtain three clusters (denoted by three colors: red, green and blue). (d) Heatmap of similarity between clusters before merging. The black solid line and the arrow mark the threshold (threshold=0.99, see Formula (2.14)). It can be seen from the heatmap, the similarity between cluster 3 and cluster 4 is above the threshold, thus merging cluster 3 and cluster 4. (e)Heatmap of similarity between clusters after merging. At this time the number of clusters is 3, and the similarity between clusters is below the threshold (threshold=0.73, see Formula (2.14)), the merging stage terminates. The number of clusters is finally determined to be 3.

To visualize the effects of the merging process, we selected the testing set C1_020 (see Evaluation) for illustration. We obtained the threshold *R_th_* in Formula (2.14) with α = 1.6. The number of clusters is 4 before merging in Figure 2b and is 3 after merging in Figure 2c. Figure 2d-e show the similarity between every two clusters measured by DBI [40] before and after merging respectively. It is worth noting that the cluster similarity between cluster 3 and 4 is above the threshold (threshold=0.99) before the merging process (Figure 2d), while all cluster similarity reaches below the threshold (threshold=0.73) after the merging (Figure 2e). Thus, our proposed algorithm can automatically determine the number of clusters through the cluster merging process.

## Evaluation

### 1. Datasets

Spike waveform data containing cluster information are generally obtained in two ways. One way is to use simulated data that quantifies algorithm performance and compares different algorithms. The other way is in-vivo extracellular recordings capturing the variability inherent in spike waveforms, which lacks in the simulated data.

#### Dataset A: simulated dataset wave_clus

In this study, we used one common simulated dataset *wave_clus* provided by Quiroga et al. [19]. In the simulation study, spike waveforms have a Poisson distribution of interspike intervals, and the noise is similar to the spikes in the power spectrum. In addition, the spike overlapping, electrode drift and explosive discharge under real conditions are simulated. To date, *wave_clus* has been used by many spike sorting algorithms for evaluating sorting performance [19–22, 24].

Dataset A contains four sets of data C1, C2, C3 and C4. Each testing set contains three distinct spike waveform templates, in which template similarity levels are significantly different (C2, C3 and C4>C1) and the background noise levels are represented in terms of their standard deviation: 0.05, 0.10, 0.15, 0.20 (C1, C2, C3 and C4), 0.25, 0.30, 0.35, 0.40 (C1). Both similarity levels and noise levels will affect the classification performance. In this study, the correlation coefficient (CC) was used to evaluate the similarity levels of spike waveforms. The higher the correlation of the two templates, the higher the similarity of the waveforms and the more difficult it will be to distinguish the two clusters.

According to spike time information, the waveforms were extracted from the *wave_clus* dataset, and then the spike alignment was conducted. Each waveform lasts about 2.5ms and is composed of 64 sample points. The peak value was aligned at the 20 th sample point.

#### Dataset B: public in-vivo real recordings HC1

HC1 is a publicly available in-vivo dataset, which contains the extracellular and intracellular signals from rat hippocampal neurons with silicon probes [44]. It is a widely used benchmark for spike sorters oriented sparse electrodes [22, 31, 38]. We used the synchronized intracellular recording as the label information of extracellular recording to obtain partial ground truth [44]. In a recent study, SpikeForest, a validation platform has evaluated the performance of ten major spike sorting toolboxes on HC1 [45].

For all the datasets, raw data were filtered by a Butterworth bandpass filter (filter frequency band 300-3000Hz), and the extracellular spikes were detected by double thresholding at Formula (2.18).

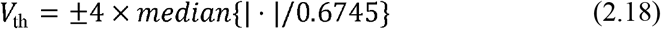

Since intracellular recording had little noise, single threshold detection was adopted to obtain intracellular action potentials. If the difference between the extracellular spike time and the intracellular peak time is within 0.3 ms, they are regarded as the same action potential. After analysis, we obtained some spikes in the extracellular recording, which corresponded to the action potentials in the intracellular recording. Thereafter we call them the marked spikes, and the rest spikes are called unmarked spikes. With regard to the typical dataset d533101 in HC1, d533101:6 contains the intracellular potential of a single neuron, while the dataset d533101:4 contains simultaneous waveforms of this single neuron as well as some other neurons. We detected 3000 extracellular spikes from extracellular recording (dataset d533101:4) and 849 intracellular action potentials from intracellular recording (dataset d533101:6). After alignment, 800 marked spikes in the extracellular recording corresponded to the action potential in the intracellular recording and were used as ground truth. The rest 2200 spikes are the unmarked spikes.

#### Dataset C:in-vivo real recordings from a non-human primate

We also compared the performance of spike sorting algorithms on in-vivo recordings from a macaque performing a center-out task in a previous study [46]. The in-vivo data were collected from a 96-channel Blackrock microelectrode array by using a commercial data acquisition system (Blackrock Microsystem, USA). Testing sets were obtained from 30 stable channels by measuring the stationarity of spike waveforms and the interspike interval (ISI) distribution.

### 2. Performance measure metrics

One of the performance measure metrics is the sorting accuracy which is the percentage of the detected spikes labeled correctly. For sample set *D*, the accuracy of classification algorithm *f* is defined as the ratio of the number of spikes correctly classified to the total number of spikes used for classification. The calculation is shown in formula (2.19):

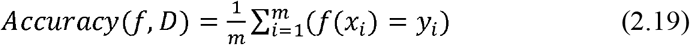

Another metric is DBI [40] that does not require prior information of clusters. DBI calculates the worst-case separation of each cluster and takes the mean value, as shown in Formula (2.20).

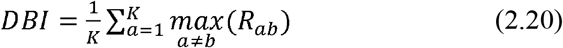

*K* denotes the number of clusters. *R* denotes the similarity between the clusters for quantitatively evaluating cluster quality. A small DBI index indicates a high quality of clustering.

To evaluate algorithm performance on real dataset HC1 with partial ground truth [45], we considered it as a binary classification problem. The classification results were divided into four cases: True Positive (TP), False Positive (FP), True Negative (TN), and False Negative (FN). We evaluated the performance of the algorithm in terms of precision rate and recall rate, as shown in Formula (2.21) and Formula (2.22), referring to the validation platform SpikeForest [45].

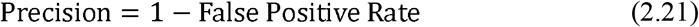

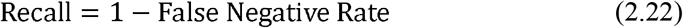

## Results

In this study, our LDA-DP algorithm was compared with five typical spike sorting methods on one simulated dataset and two real datasets concerning several performance measure metrics. For comparison, we choose the algorithm LDA-GMM [22], which has the same feature extraction method as LDA-DP, and choose the algorithm PCA-DP, which has the same clustering method. Then we select two classic and widely-used spike sorting algorithms, PCA-Km [13] and LE-Km [27], along with a recently proposed algorithm GMMsort [31]. SpikeForest has compared the performance of ten spike sorting methods on several neural datasets from both high-density probes (like neuropixel) and sparse probes (like classical tetrodes or microwire arrays). IronClust outperforms the other 9 methods on Dataset B [47]. Thus we quoted these results and compared them with our LDA-DP and the above-mentioned 5 algorithms. These algorithms are all unsupervised and automated, except that GMMsort needs some manual operation in the last step of clustering. In comparison, the feature subspace dimension was fixed as 5 for GMMsort [31] by default and 3 for the rest algorithms [13,22,27].

### 1. Performance comparison in the simulated Dataset A

A prominent feature extraction method can find the low-dimensional feature subspace with a high degree of differentiation, which is the basis of the high performance of the whole algorithm. Thus, we compared the robustness of different feature extraction methods.

As the noise level or waveform similarity increases, the feature points of different clusters will gradually get closer in the feature subspace, the inter-class distance will decrease, and the boundary will be blurred, increasing classification difficulty. Thus, we chose the testing set C3 with high waveform similarity as the testing set to compare the performance and the noise resistance for five feature extraction methods (PCA-KM and PCA-DP used the same feature extraction method PCA).

When the noise level increases, the standard deviation of each waveform template increases (Waveforms column in Figure 3), bringing difficulties to feature extraction. In this case, feature points extracted by the LDA method in LDA-DP and LDA-GMM are clustered separately, while in the contrast, feature points from the rest three methods are overlapped to some degree. Even under the worst condition when the noise level rises to 0.20, the proposed LDA-DP algorithm has the least overlapped feature points among the five methods. It notes that the feature extracted by the LDA-DP algorithm has high robustness to noise and waveform similarity.

**Figure 3.**
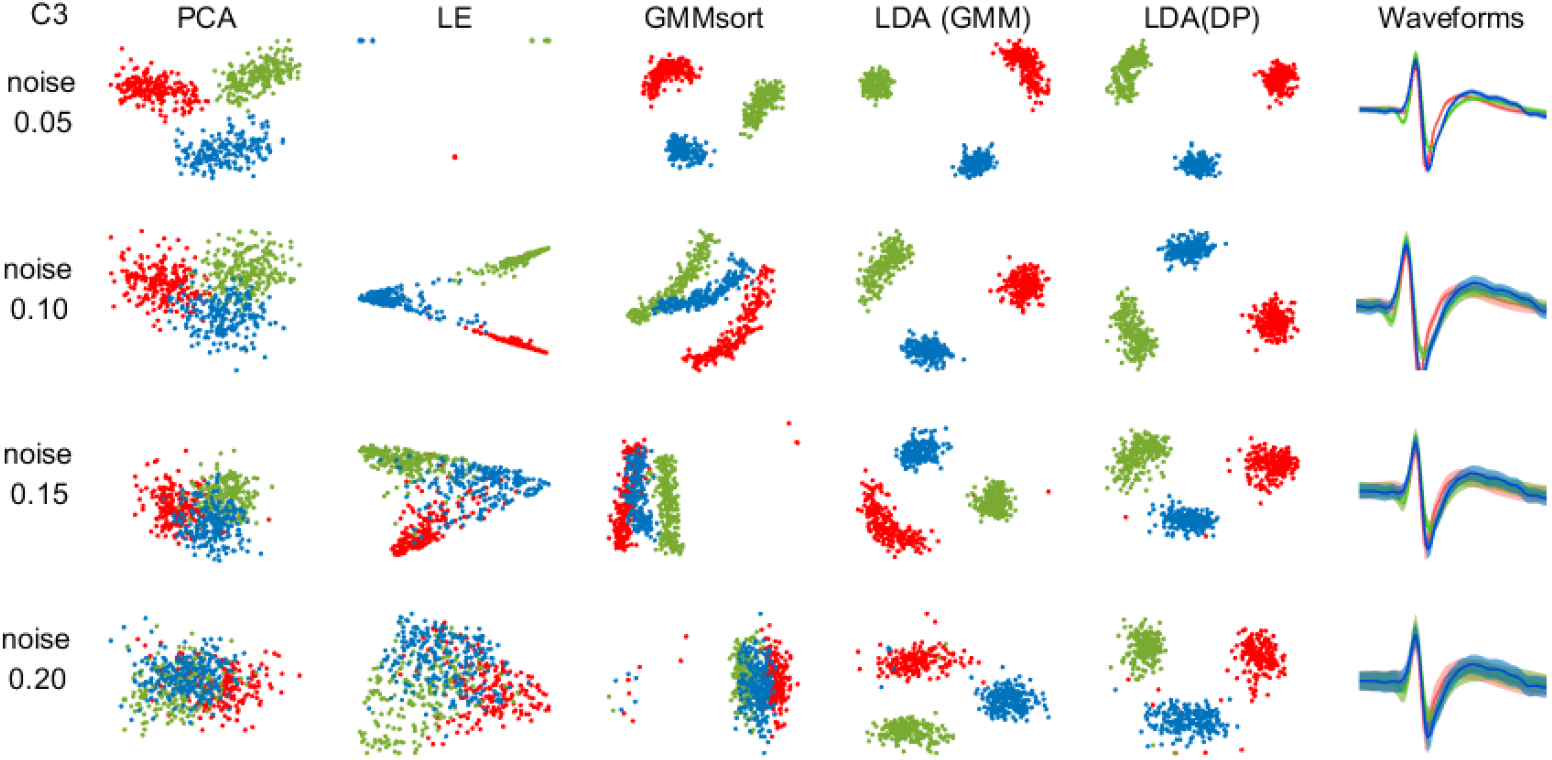
Two-dimensional feature subspace for algorithms on dataset C3 under four noise levels. For comparing the feature extraction capability of each algorithm, the data points were colored according to the ground truth. Red, green and blue dots represent the features extracted from three clusters, respectively. The last column shows the average spike waveforms of three clusters obtained by LDA-DP, and the shaded part represents the standard deviation.

We examined the performance of the 6 algorithms (PCA-Km, LE-Km, PCA-DP, LDA-GMM, GMMsort, and LDA-DP) on the Dataset A, excluding the overlapping spikes. In order to compare the robustness of each algorithm, two performance metrics were employed in this study: sorting accuracy and cluster quality. For each testing set, 5-fold cross-validation was performed. For PCA-Km and LE-Km, the number of clusters was set to 3; And for PCA-DP, LDA-GMM, GMMsort, and LDA-DP, the number of clusters can be determined automatically.

Table 1 presents the average and the standard deviation (std) of the sorting accuracy. It is worth noting that the average sorting accuracy of LDA-DP on most of the testing sets is higher than that of the other methods. At the same time, LDA-DP also achieves a lower standard deviation of the average accuracy on most of the testing sets.

**Table 1.**
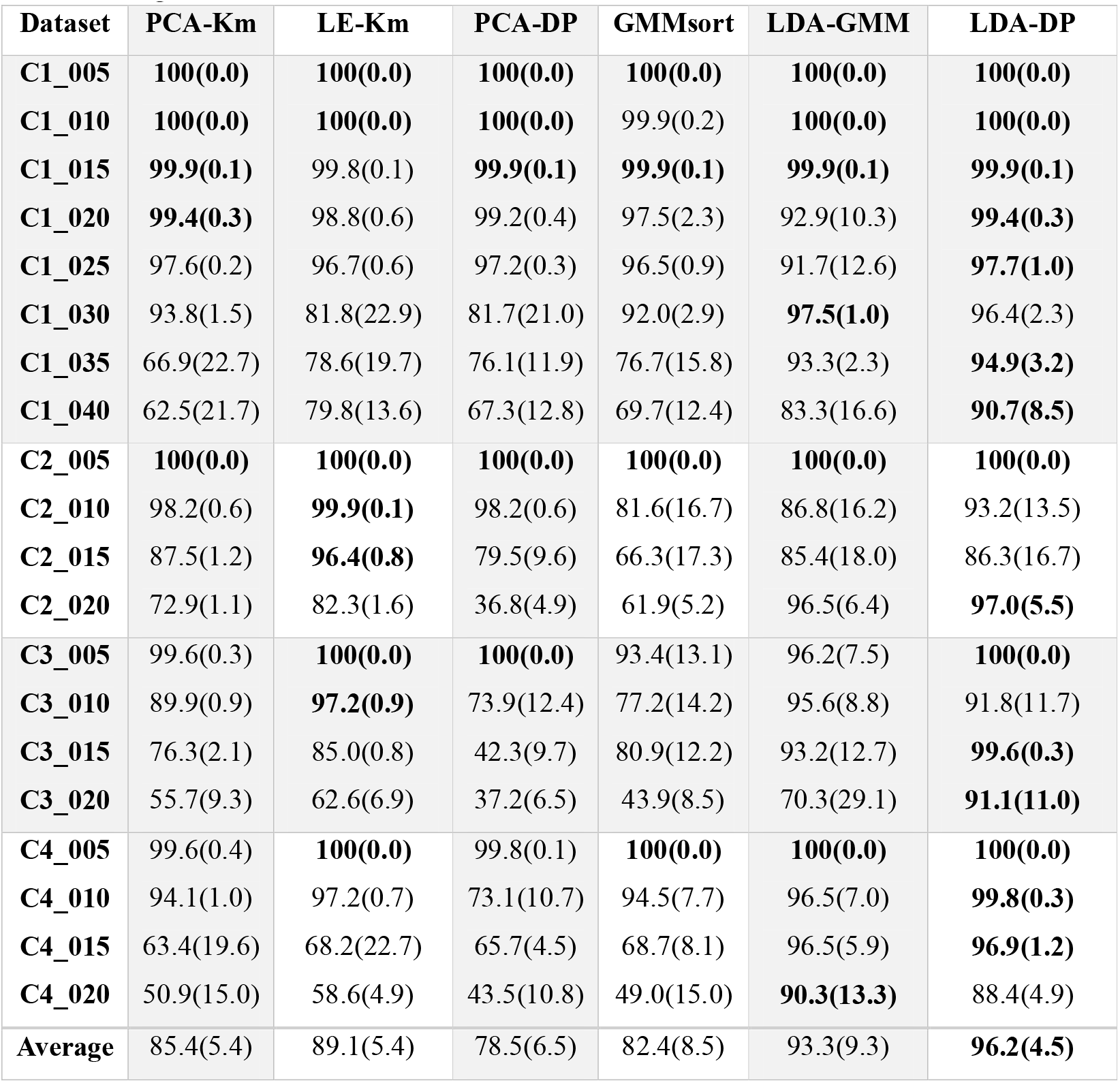
Average sorting accuracy percentage on the Dataset A (simulated dataset *wave_clus*) (with no overlapping spikes) for 6 algorithms. The standard deviation of the accuracy is in parenthesis. The bold number represents the best performance in each testing set.

In the testing set C1 whose classification difficulty is low, most algorithms achieve high accuracy. As the noise level rises to 0.40, the sorting accuracy of the rest five algorithms drops below 90%, but the accuracy of LDA-DP is still up to 90.7%. Moreover, in the testing set C2, C3 and C4, when both the waveform similarity and the noise level increase, only LDA-DP and LDA-GMM can maintain high accuracy relatively. The comparative results indicate that these two algorithms have a great power to distinguish waveforms and are highly resistant to noise. LDA-DP is especially outstanding because it maintains a higher sorting accuracy steadily (>85%). On average, the mean accuracy of LDA-DP reaches 96.2%, which is the highest in all 6 algorithms. At the same time, LDA-DP achieves the lowest mean standard deviation (std=4.5).

To further visualize the robustness of each algorithm concerning noise, we plotted the changing curve of performance with four or eight noise levels on 4 simulated datasets (C1, C2, C3 and C4). Figure 4a, c, e and g show the accuracy curve, while Figure 4b, d, f and h show the DBI curve. In Figure 4, as the noise level increases, the performance of all algorithms drops (The sorting accuracy decreases and the DBIs increase). When the noise level is low, all of the 6 algorithms get high accuracy and low DBI. The gaps between algorithms are not obvious. However, when the noise levels increase, the performance of PCA-Km, LE-Km, PCA-DP and GMMsort deteriorates. And in most cases, LDA-DP performs better than LDA-GMM. In all simulated data, LDA-DP displays a high level of performance: the sorting accuracy rate is above 85%, and the DBI is below 1.5, which is generally superior to other algorithms and shows high robustness to noise.

**Figure 4.**
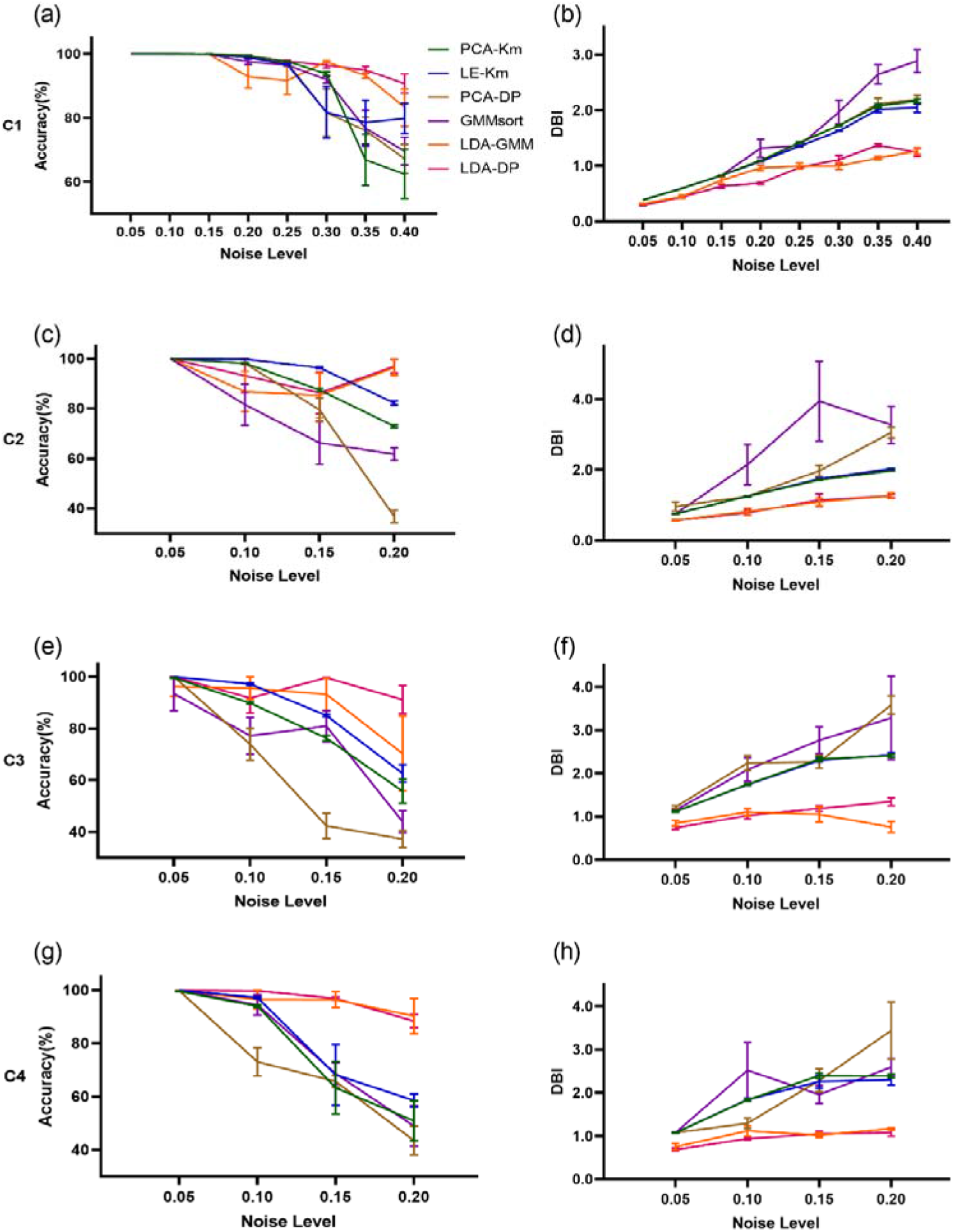
The sorting performance of 6 spike sorting algorithms on the Dataset A concerning noise levels. (a) We compared sorting accuracy of PCA-Km, LE-Km, PCA-DP, LDA-GMM, GMMsort and LDA-DP with respect to noise on the testing set C1 (noise level: 0.05, 0.1, 0.15, 0.2, 0.25, 0.3, 0.35, 0.4). Error bars represent s.e.m.. (b) DBI comparison concerning noise on the testing set C1. The evaluation was performed as in a. (c)-(h) Sorting accuracy and DBI concerning noise on the testing set C2, C3 and C4 respectively.

We also compared the robustness concerning waveform similarity. In the right side of Figure 5a, the shapes of the three waveform templates were plotted for four testing sets. The correlation coefficients (CC) of the three templates were used to measure the similarity level in each testing set (Figure 5a left). The results indicate that the waveform similarity in C2, C3 and C4 is significantly higher than that in C1 (Paired t test, p<0.01), thus the classification of C2, C3 and C4 is relatively more difficult. In order to intuitively show the algorithm performance differences, we chose to plot the accuracy and DBIs of 6 algorithms on the four testing sets: C1_020, C2_020, C3_020 and C4_020, in which the waveform similarity is diverse and the noise level remains the same (Figure5b-c). In Figure 5b, LDA-DP is superior to the other algorithms in terms of sorting accuracy. In the case of high waveform similarity, the accuracy of other algorithms fluctuates somewhat, while the accuracy of LDA-DP still maintained at a high level. For the DBIs (Figure 5c), the cluster quality of LDA-DP is also promising.

**Figure 5.**
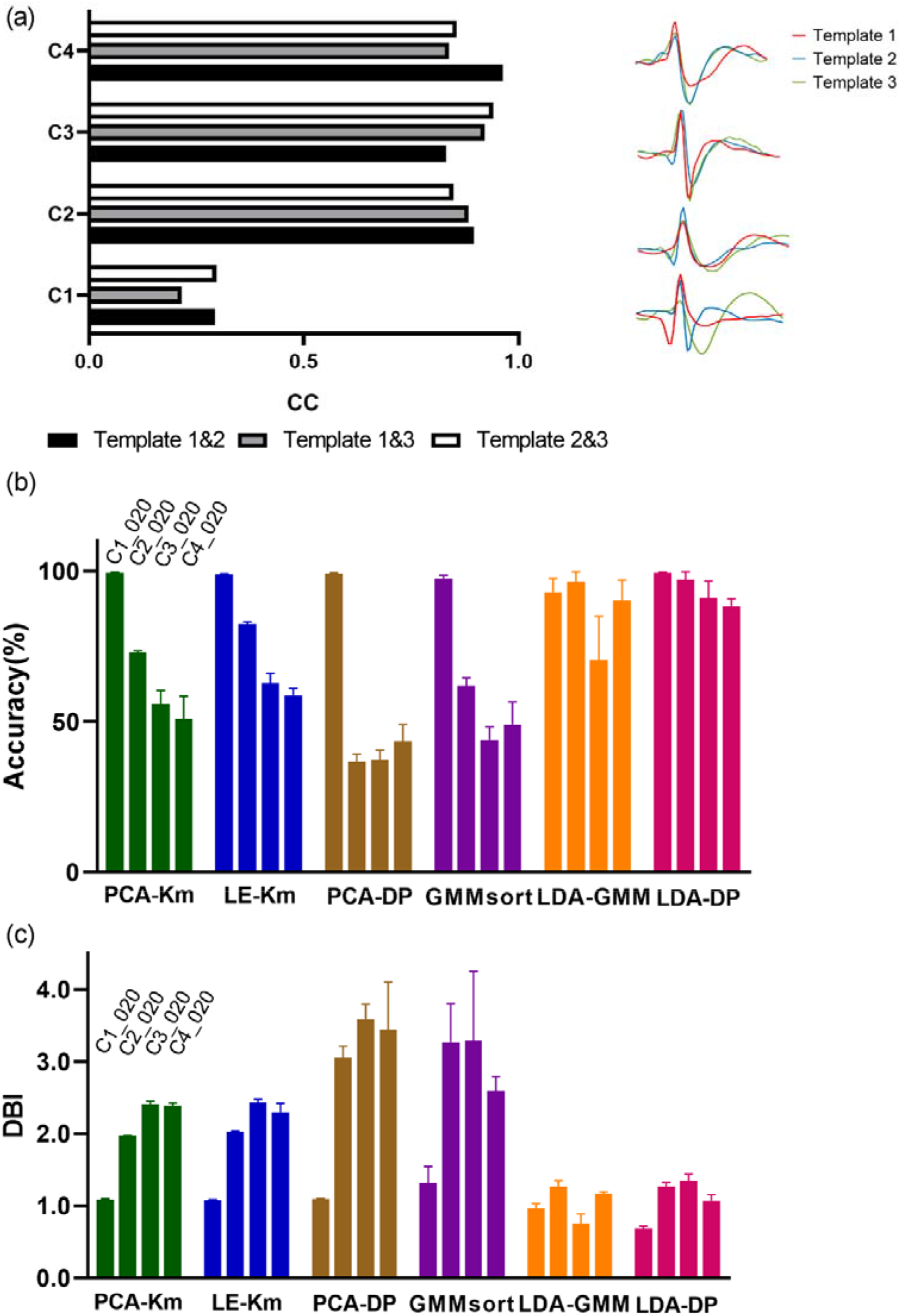
The sorting performance of 6 spike sorting algorithms concerning waveform similarity using the testing set C1_020, C2_020, C3_020 and C4_020. (a) Comparing the similarity of waveform templates in four testing sets. Left: bar graph of the correlation coefficient (CC) of the three waveform templates for four datasets. Black, gray and white respectively represent CC of template 1 and template 2, CC of template 1 and template 3, and CC of template 2 and template 3. Right: Waveform templates of each testing set. Red, blue and green represent template 1, template 2 and template 3, respectively. (b) Bar graph of sorting accuracy of 6 spike sorting algorithms on the testing set C1_020, C2_020, C3_020 and C4_020. Error bars represent s.e.m. (c) Bar graph of DBI of 6 spike sorting algorithms on the testing set C1_020, C2_020, C3_020 and C4_020. Error bars represent s.e.m.

### 2. Performance comparison in the in-vivo Dataset B

To further evaluate the performance of our algorithm on in-vivo datasets, we compared the performance of LDA-DP and the above 5 algorithms on Dataset B. For PCA-Km and LE-Km, the number of clusters was manually set to 3; And for PCA-DP, LDA-GMM, GMMsort, and LDA-DP, the number of clusters can be determined automatically. Firstly, one dataset d533101 in Dataset B, which was widely adopted in previous studies [22, 31, 38], was chosen for illustration. Figure 6a shows the two-dimensional feature subspace extracted by each method. Data points are grouped into three clusters in the subspace and the waveform panel shows the average spike waveforms of the three clusters obtained by LDA-DP. The figure suggests that the LDA method successfully extracts optimal feature subspace, benefiting from the credible feedback of the clustering method (DP) through several iterations. The three clusters are much more distinct in the subspace of LDA (GMM) and LDA (DP) than in other methods.

**Figure 6.**
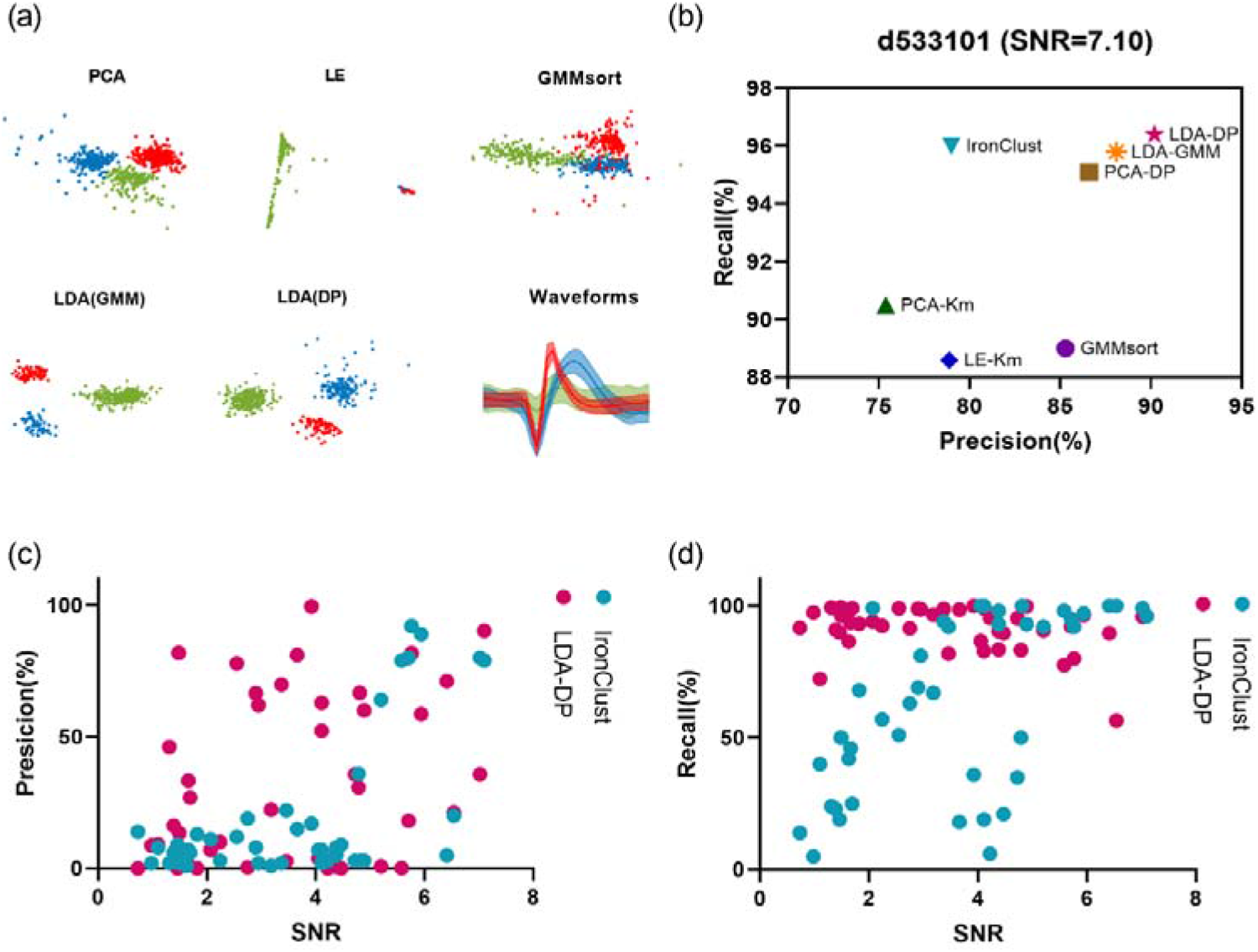
The sorting results on the in-vivo Dataset B when comparing 6 spike sorting algorithms. (a) Two-dimensional feature subspace extracted by each method. The data points are colored according to the sorting results, in which red, green and blue represent cluster 1, cluster 2 and cluster 3 respectively. The waveform panel shows the average spike waveforms of three clusters obtained by LDA-DP, and the shaded part represents the standard deviation. (b) The precision rate and the recall rate obtained according to partial ground truth on dataset d533101 in HC1. (c-d) The precision rate and recall rate comparison with regard to SNR on all the 43 datasets in Dataset B.

According to the partial ground truth, we analyzed the classification results of algorithms by evaluating the precision rate and the recall rate on the d533101 (SNR=7.10). Figure 6b shows the classification results of 6 algorithms, along with one outstanding spike sorting software package, IronClust [47], which is benchmarked by the SpikeForest [45]. Compared with other methods, LDA-DP has a maximum precision rate, as well as a maximum recall rate. Although IronClust is a density-based sorter and shows top average precision and recall rate among the 10 toolboxes validated in SpikeForest, its precision rate on the sparse recording is inferior to some algorithms developed for sparse neural recording.

For further performance evaluation, we compared LDA-DP and IronClust on all the 43 datasets in Dataset B, with regard to signal-to-noise ratio (SNR). In Figure 6c, LDA-DP has the higher precision rate on 25 datasets among all 43 datasets. Moreover, under a lower SNR (SNR<4.0), the precision of LDA-DP is higher on 17 datasets out of 23 datasets. In Figure 6d, LDA-DP has the higher recall rate on 28 datasets among all 43 datasets. Especially, under the lower SNR (SNR<4.0), the recall rate of LDA-DP is higher on 21 datasets out of 23 datasets, demonstrating robustness convincingly. Our results indicate that LDA-DP also outperforms other algorithms on the Dataset B.

### 3. Performance comparison in the in-vivo Dataset C

In order to test the robustness of the algorithm in the real case with more complex distribution characteristics, we also compared the performance of 6 algorithms on the in-vivo Dataset C. In particular, as shown in Figure 7a, spikes from one typical channel 54 in dataset C present a more messy distribution in the two-dimensional feature subspaces extracted by each method. The data points are colored by the sorting results and are grouped into four clusters. The Waveforms column shows the shapes of the four spikes due to the sorting results of LDA-DP. It indicates that these four spikes have highly similar shapes. The above complication may pose huge challenges to feature extraction. As Figure 7a suggests, features extracted by LDA(DP) are apparently more separable than all other methods, leading the subsequent clustering to be more accurate. We notice that the data points show nonspherical-distributed in the LE subspace. Since clustering methods, such as K-means, have poor performance in identifying the clusters of nonspherical distribution, LE-Km may encounter difficulties in clustering features. We further compared the cluster quality of 6 algorithms using the spikes in this channel. For PCA-Km and LE-Km, the number of clusters was manually set to 4; And for PCA-DP, LDA-GMM, GMMsort, and LDA-DP, the number of clusters can be determined automatically. In Figure 7b, the DBI index of LDA-DP is significantly lower than those of other algorithms (* p<0.05, ** p<0.01, Kruskal-Wallis test), indicating that LDA has higher cluster quality and better performance than other algorithms. Moreover, we conducted a comparison on all 30 channels in Dataset C. The results are shown in Figure 7c, the median of the DBI index for the LDA-DP is lower, and in general, LDA-DP has a significantly higher cluster quality (* p<0.05, *** p<0.001, Kruskal-Wallis test). Thus, LDA-DP also demonstrates outstanding robustness advantages on Dataset C, which is consistent with the results from the previous two datasets.

**Figure 7.**
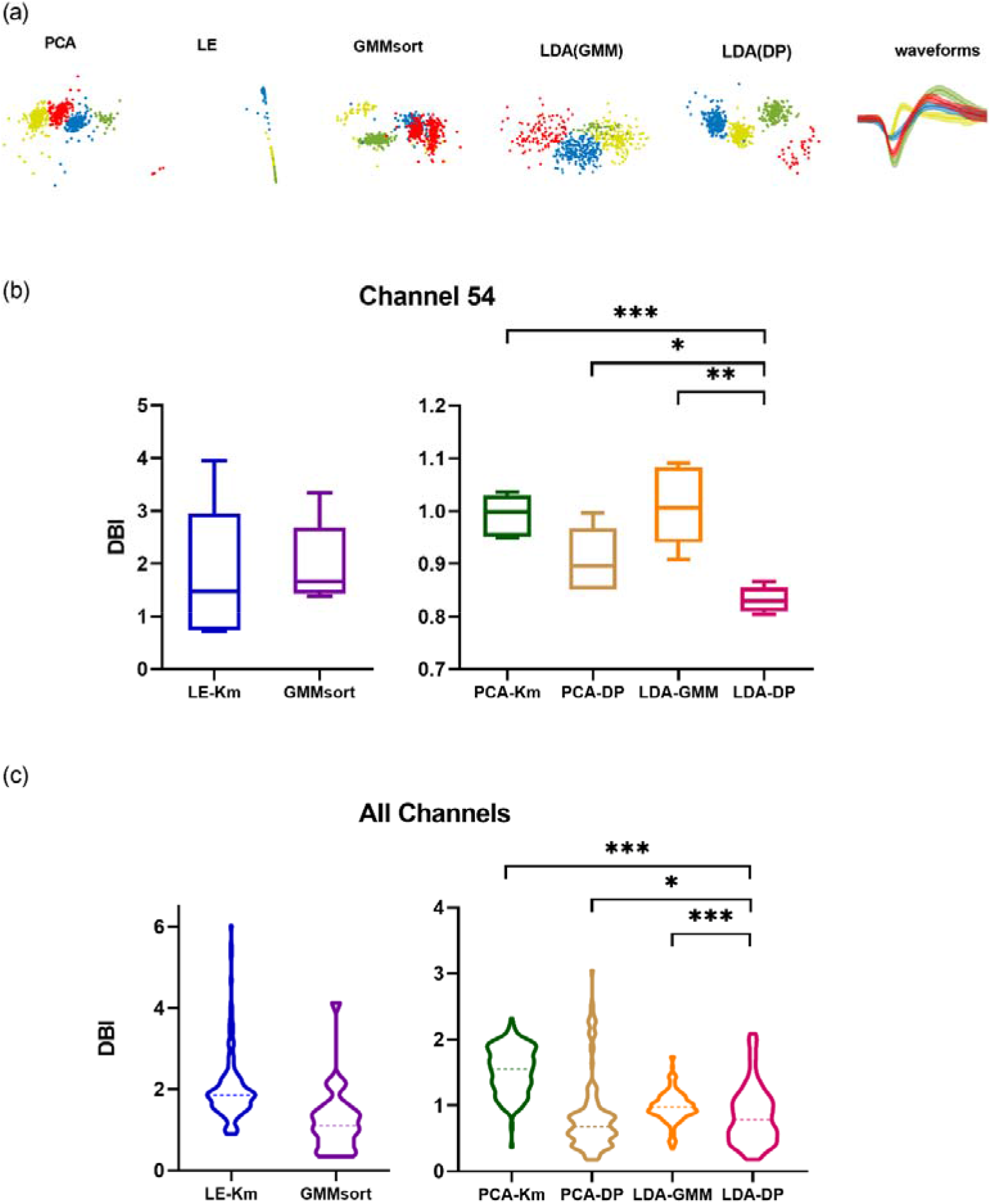
The sorting results on Dataset C when comparing 6 spike sorting algorithms. (a) Two-dimensional feature subspace extracted by each method. The data points are colored according to the sorting results, in which red, green, yellow and blue represent cluster 1, cluster 2, cluster 3 and cluster 4 respectively. The last column shows the average spike waveforms of four clusters obtained by LDA-DP, and the shaded part represents the standard deviation. (b) Comparison of cluster quality using spikes from one particular channel 54 in Dataset C. We conducted a 5-fold cross-validation and draw boxplots of the DBI index for 6 algorithms. * p<0.05, ** p<0.01, *** p<0.001, Kruskal-Wallis test. (c) Comparison of cluster quality using spikes from all 30 channels in Dataset C. We conducted a 5-fold cross-validation for each channel and then drew the violin plot. The dashed lines represent the median of the DBI. * p<0.05, *** p<0.001, Kruskal-Wallis test.

## Discussion

In this study, our proposed LDA-DP competed with six algorithms on both simulated and real datasets. The LDA-DP exhibits high robustness on both simulated and real datasets. For the simulated dataset wave_clus [19], the LDA-DP maintains an outstanding sorting accuracy and cluster quality, indicating high robustness to noise and waveform similarity. And on the real dataset HC1 [44], the comparison further illustrates the robustness of LDA-DP under low SNR. We finally evaluated the algorithm on real data from non-human primates [46]. The performance of LDA-DP also exceeds other algorithms when facing more complex data distributions.

In this study, the performance of LDA-DP and LDA-GMM is significantly better than the other 4 algorithms (Figure 4-7). We can see the gap between LDAs and non-LDAs, for example, LDA-DP and PCA-DP. It is probably because the LDAs are supervised methods while the other methods are unsupervised. Through multiple iterations, LDA finds the optimal feature subspace based on the feedback provided by the clustering method, while unsupervised methods get no feedback. Therefore, the advantages of LDA-DP and LDA-GMM benefit from the combination of the feature extraction method LDA and the cluster method (DP or GMM). Our results are consistent with previous studies [22, 24].

Although two LDAs have high performance, LDA-DP has a better performance than LDA-GMM (Figure 6-7) on in-vivo datasets. Due to the joint optimization framework, the feature extraction method LDA also benefits from an outstanding clustering method. Since DP does not make any assumption about data distribution, the DP [39] method has more advantages when processing real data with more complex characteristics. GMM [48], as its name implies, assumes that data follows the Mixture Gaussian Distribution. This sort of fitting often encounters difficulties in dealing with more complex situations, where real data are not perfectly Gaussian distributed. Several studies in other fields have encountered similar problems [49–53].

Additionally, some classic clustering methods, such as K-means, specify the cluster centers and then assign each point to the nearest cluster center [28]. Thus, this kind of methods perform poorly when applied to nonspherical data. In the contrast, the DP algorithm is based on the assumption that the cluster center is surrounded by points with lower density than it, and the cluster centers are relatively far apart. According to this assumption, DP identifies cluster centers and assigns cluster labels to the rest points. Therefore, it can well deal with the distribution of nonspherical data. This is one of the potential reasons for the outstanding performance of LDA-DP.

It is worth noting that LDA-DP is an automated algorithm. Although the values of some parameters may affect the final results, we can still preset some optimized values to avoid manual intervention during operation. For example, in this study, we fixed the threshold coefficient *α* as 1.6 and verified its high performance on one simulation dataset and two in-vivo real datasets. Although whether this optimal value fits all datasets needs to be tested and evaluated with more data, the current evaluation has fully demonstrated the general applicability of this value.

Most methods compared in our study were employed to sort the spike data from a single microelectrode. These spike data can be collected through sparse probes such as Utah arrays and microwire arrays that are widely used in neuroscience and have advantages in the stability of long-term recording. Thus our algorithm is not comparable with spike sorters for high-density probes and cannot be verified on the data from high-density electrodes. In the future, we will try to swap out the clustering steps in high-density spike sorters such as Kilosort [54] or Mountainsort [55] with DP to see whether the DP would bring an accuracy improvement in the neural recordings from high-density probes.

## Conclusion

By combining LDA and DP to construct a joint optimization framework, we proposed an automated spike sorting algorithm and found it is highly robust to noise. Based on the iteration of LDA and DP, the algorithm makes the feature extraction and clustering benefit from each other, continuously improving the differentiation of feature subspace and finally achieves high spike sorting performance. After evaluation on both simulated and in-vivo datasets, we demonstrate that the LDA-DP meets the requirements of high robustness for sparse spikes in the cortical recordings.

## Data Availability

The code of the LDA-DP used in the current study is available at https://github.com/EveyZhang/LDA-DP.

## Notes

### Competing Interest Statement

The authors have declared no competing interest.

